# Unifying regulatory motifs in endocrine circuits

**DOI:** 10.1101/2024.08.06.606903

**Authors:** Moriya Raz, David S. Glass, Tomer Milo, Yael Korem Kohanim, Omer Karin, Avichai Tendler, Avi Mayo, Uri Alon

## Abstract

In this study we identify unifying design principles in human endocrine systems. We find that 43 hormone systems, controlling diverse physiological functions, fall into 5 classes of circuits with shared structure – thus only a small number of the possible circuits actually occur. Each class uses a different regulatory logic to perform specific dynamical functions, such as homeostasis, acute input-output response or adjustable set points. The circuits employ interactions on two timescales: hormone secretion on the scale of minutes-hours and growth and shrinkage of endocrine gland mass on the scale of months, which impacts the amount of hormone the glands secrete. This two-timescale principle recurs in several classes of circuits, including the most complex class, which has an intermediate gland, the pituitary. We analyze the pituitary circuit in detail and find tradeoffs between endocrine amplification, buffering of hypersecreting tumors, and rapid response times. These unifying principles of regulation build a foundation for systems endocrinology.

## Introduction

Hormones are regulatory molecules secreted by endocrine cells into the circulation, where they affect target tissues(1). Hormones impact physiological functions including growth, metabolism, reproduction, stress response, immunity, mood and behavior(1–6). Dysregulation of hormones underlies a wide range of pathologies including diabetes, thyroid disorders, fertility disorders, growth delay, Cushing’s syndrome, anemia and mood disorders(1,7).

Hormones share a common aim – to regulate physiological function. However, they differ in their regulatory logic (1). Some hormones are secreted under the control of neuronal stimulation, such as antidiuretic hormone (ADH) (8). Some are secreted under the control of other hormones, such as thyroxine, which is controlled by thyroid-stimulating hormone (TSH) (9). Others are secreted in response to metabolic signals, such as insulin, which is secreted in response to elevated blood glucose(10). Yet other hormones are regulated in complex cascades. For example, several key hormones regulated by the hypothalamus go through the pituitary, an intermediate gland that controls multiple effector glands(11).

We ask what determines the regulatory logic for a given hormone. In order to do so, it is useful to define recurring patterns and concepts(12). This systems-level approach was successful in deriving general principles in gene regulation networks and other complex biochemical systems(13). Well-known principles in endocrinology likewise have been instrumental in guiding experimental and clinical studies. For example, feedback regulation is a hallmark of homeostatic endocrine systems such as insulin control of glucose(14). The discovery of such feedback loops led to mathematical models that resulted in formulas that are useful for clinical research – such as estimates for insulin resistance known as the HOMA-IR formula(15,16). Furthermore, analogies between systems allow researchers to carry over ideas from one system to another, such as the development of HOMA-like formula for the thyroid(17,18), and the discovery of long feedback loops in the hypothalamic-pituitary hormone axes(19,20).

Recent work has defined regulatory principles related to changes in endocrine gland mass, under control of the hormones in the same pathway(21–23). This allows gland mass to grow or shrink, and thus to produce more or less hormone. For example, the thyroid grows when iodine is scarce in order to produce more thyroxine. Mass changes on the scale of weeks compensate for physiological changes such as insulin resistance(24), pregnancy(25), addiction(26). They govern responses over months to chronic stress(21), hormone seasonality(23) and the transition between subclinical and clinical autoimmune disease in which upstream glands can partially compensate for destruction of tissue(22).

There remain open questions which we aim to address here. What determines the regulatory logic of each hormone circuit? How does the regulatory logic affect the function of the circuit? Why doesn’t the hypothalamus directly communicate with target organs like the adrenal, and instead goes through an intermediate gland – the pituitary? The answers to these questions can deepen our understanding of the endocrine system and contribute to the field of systems endocrinology.

In this study we identify unifying design principles in the human endocrine system. In order to do this, we studied 43 hormone systems, seeking to categorize them based on regulation circuitry. We find that diverse systems can be organized into 5 classes of circuits according to shared regulatory motifs – regardless of biochemical detail. These classes are only a tiny fraction of the possible circuits. Using dynamical mathematical models of circuit function, we show that each class offers specific regulatory functions. We show how the pituitary gland, an intermediate gland – a key element of several endocrine circuits – can offer functional advantages over alternative designs that lack such an intermediate gland. These include amplification of hormone secretion, buffering against hormone-secreting tumors and providing speedup of response to chronic stress. These principles enhance our understanding of human hormone circuits.

## Results

### Hormone circuits can be categorized into five distinct classes

There are 64 known hormones in humans(4). We analyze the main regulatory interactions as described in the primary textbooks in clinical endocrinology(1) and complemented this by a literature search. For each hormone, we identified the main regulatory interactions that control its secretion, the gland or tissue that secretes the hormone and the major physiological variable (metabolite or other hormone) that is affected by the hormone. We call this set of interactions a hormone circuit. These circuits represent baseline biological consensus under normal physiological conditions. We found sufficient information to systematically categorize the regulation of 43 of these hormones (Methods). Examples of hormones not included due to lack of information include apelin and neurokinin B (full list in SI).

We found that the hormone systems that we studied can be organized into five classes of circuits, which we term classes 1-5 (Figure 1). The various hormone systems within each class obey the same circuit logic. By analogy with gene-regulatory and other networks, we describe these classes as circuit motifs(13). Note that these circuits differ from classical gene-regulatory motifs that act within the cell because they include cell-cell signaling and cellular growth. Class 1 circuits are the simplest and class 5, which have three glands, are the most complex.

**Figure 1:**
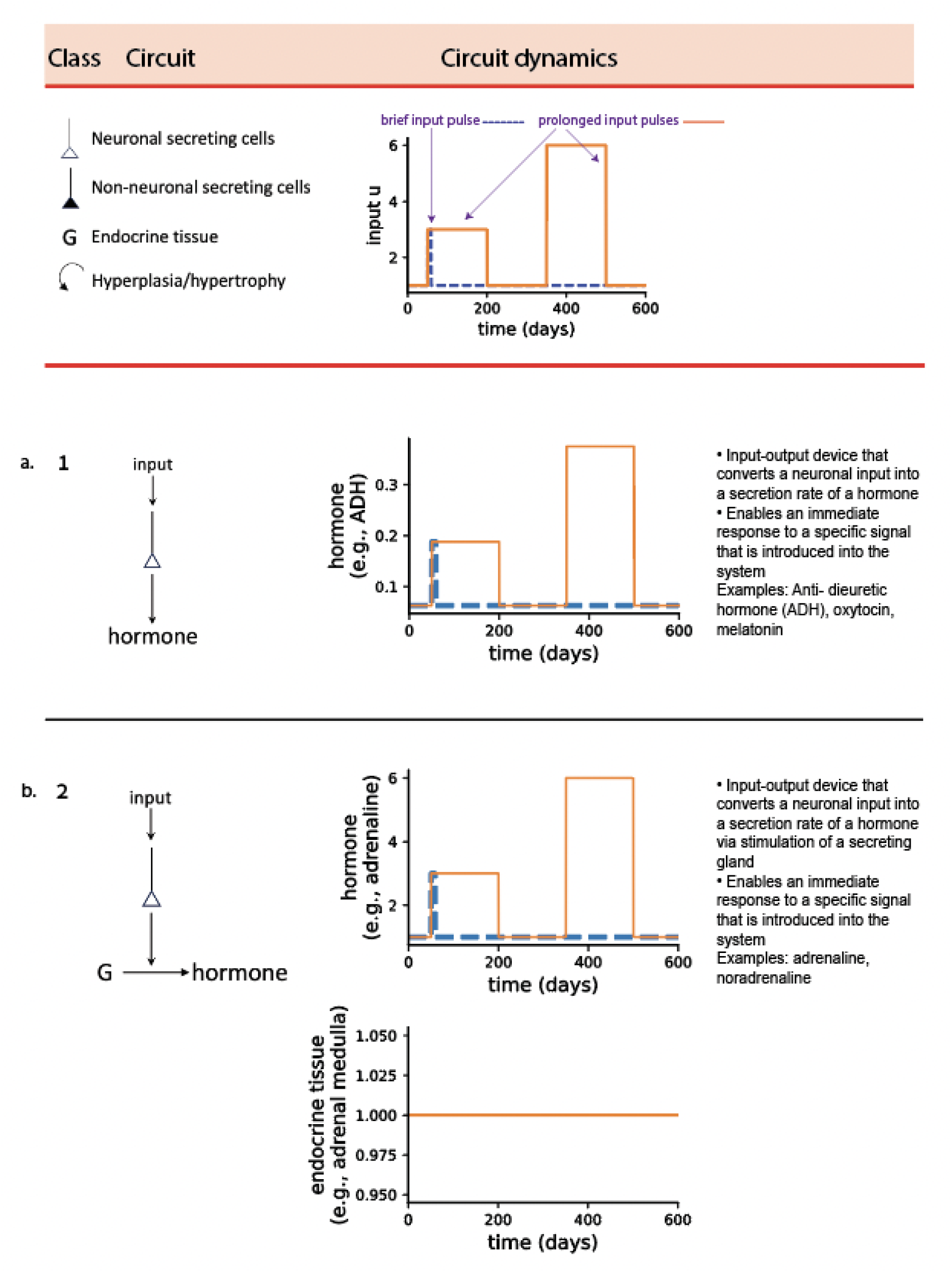
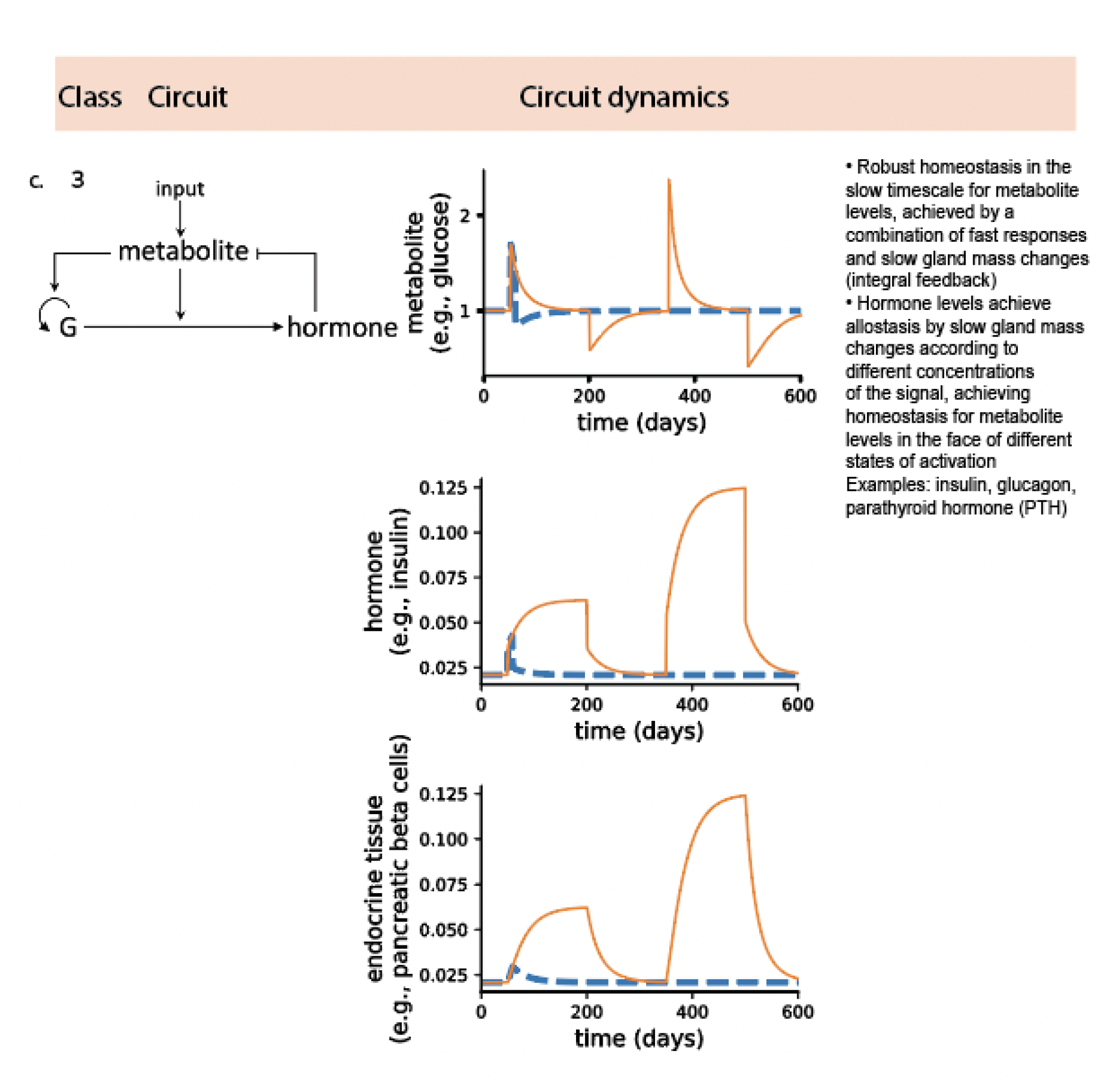
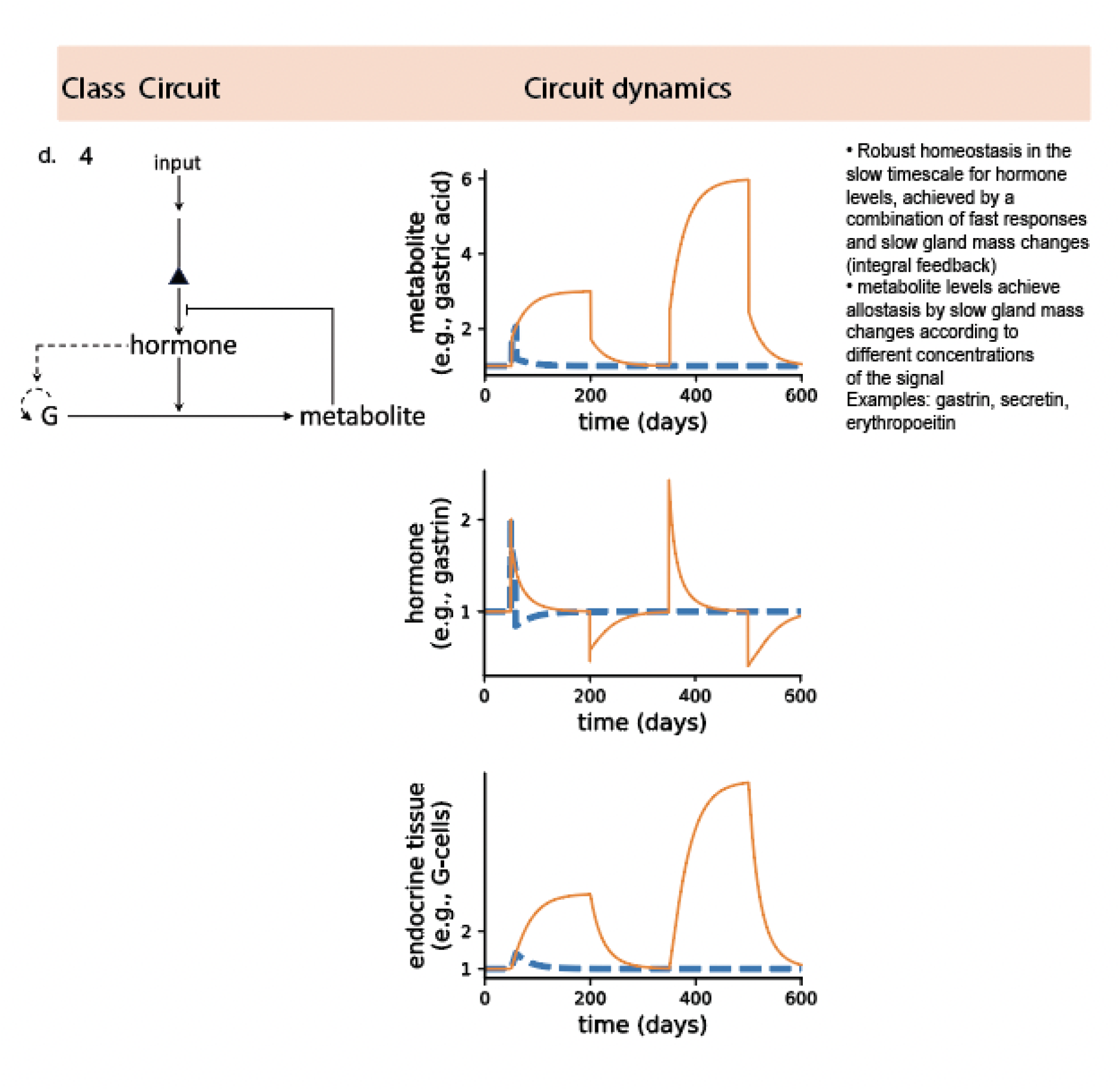
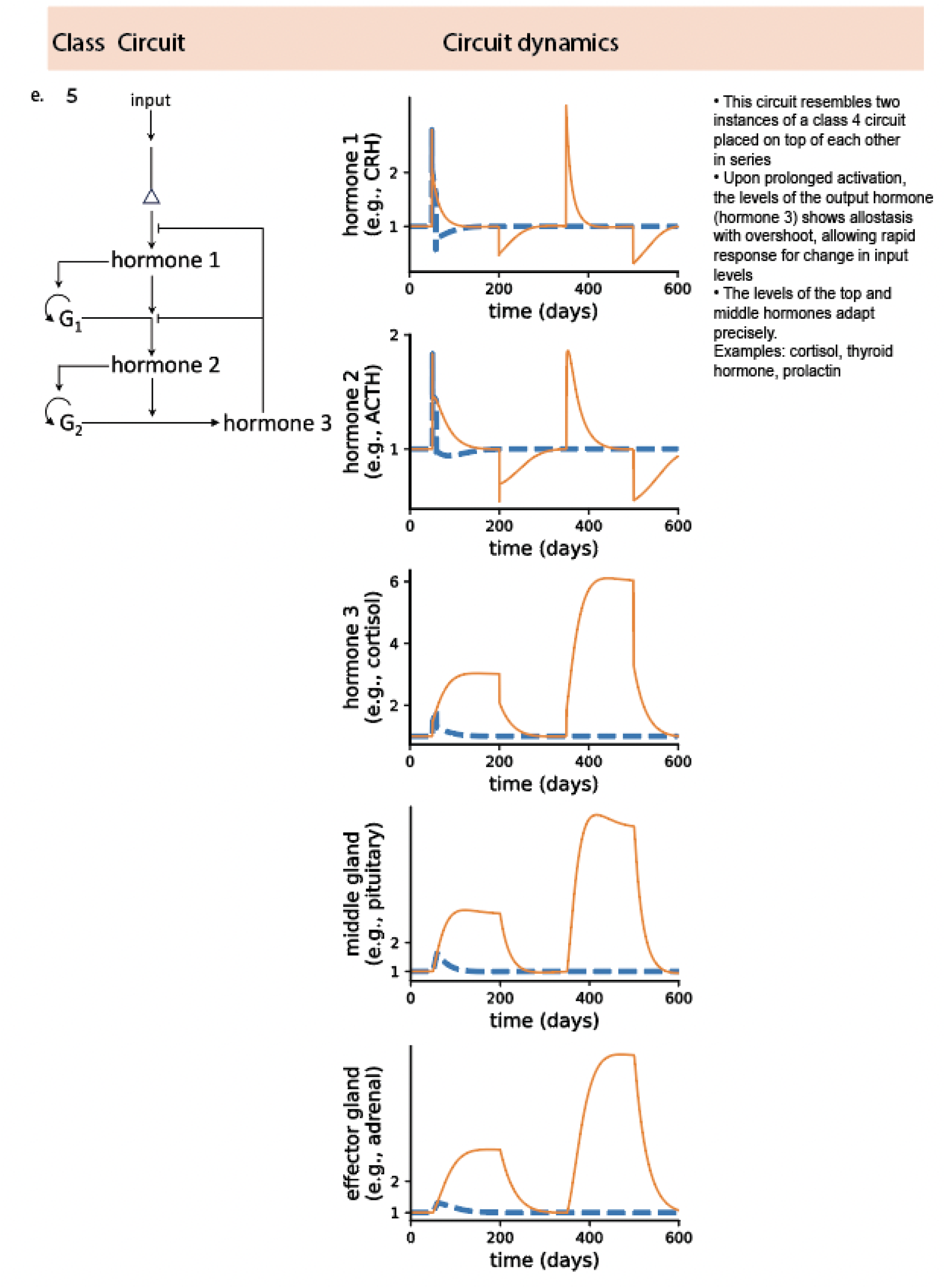
The five classes of endocrine circuits and their dynamic response to brief and prolonged input stimuli. **(a-e)** Class 1 – 5, respectively. Metabolite, hormone and gland mass dynamics from simulations with a brief input pulse of 9 days (dashed blue line) or a prolonged pulse of 150 days (orange line) and a second prolonged pulse with twice the amplitude at day 350. The functions of the circuits are summarized. Simulation parameters are *q* = *q*_3_ = 16 *day*^−1^, *q*_1_ = 288 *day*^−1^, *q*_2_ = 48 *day*^−1^, *r*_*Classes* 1−3_ = *r*_3_ = 16 *day*^−1^, *r* = 288 *day*^−1^, *r*_*Class* 4_ = *r*_2_ = 48 *day*^−1^, *c* = *c*_1_ = *c*_2_ = *d* = *d*_1_ = *d*_2_ = 1/15 *day*^−1^, *s* = 48 *day*^−1^, *a* = 16 *day*^−1^ (Box 1, Methods).

We note that the number of possible regulatory circuits is much larger than 5. For example, if one enumerates all possible connected circuits of three glands one obtains 512 possible circuits (6 interactions between the glands and 3 possible autocrine regulations providing a total of nine interactions, each of which can be present or absent totaling 2^9^ = 512 circuits). If one includes all combinations of possible regulatory signs on the arrows (each either negative or positive), this number grows to 3^9^ = 19683 possibilities. Thus it appears that physiology utilizes only a tiny fraction (0.025% or 1%) of the possibilities, suggesting a selected, meaningful function for the 5 classes.

For each class of circuits we developed a simple mathematical model (Box 1). These models describe the dynamics of the hormone concentration, metabolite concentrations and the gland functional mass. The models are based on previous approaches – the Topp model for the insulin system(27) and the mathematical model for the Hypothalamic-Pituitary-Adrenal (HPA) axis by Karin et al (21). The models have a fast (hours) timescale of hormone production and removal, and a slow (weeks) timescale of gland mass changes.

The models describe the essential mechanisms with a minimal number of parameters (production rates, removal rates, growth rates). We constructed a model for each of the circuit classes, and used it to study the circuit dynamics and function. We provide simulations of the dynamics of the hormones and metabolites for each type of circuit, for brief and prolonged stimulations (Figure 1). For brief stimulation we chose a 9 day pulse, much shorter than the typical timescale of gland-mass changes, and for a prolonged stressor input we used 150 days, much longer than the gland change timescale. We then provided a second input pulse of twice the amplitude. In each case, the simulations indicated distinct behavior related to the circuit topology. We discuss each of these behaviors and their sensitivity to parameters.

### Class 1 and 2 circuits function as fast input-output devices

Class 1 and class 2 circuits, the simplest classes, are both input-output devices that convert a neuronal input into a secretion rate of a hormone. Class 1 circuits (Figure 1a) are simply neurons that directly secrete a hormone. An example is the system of hypothalamic neurons that secrete ADH and oxytocin(8). Class 2 circuits (Figure 1b) have a neuronal input to an endocrine cell. Examples include sympathetic control of adrenal medulla cells that secrete adrenaline and noradrenaline(28,29). Simulations of class 1 and 2 circuits responded to an input pulse on the fast timescale of minutes to hours. The response was elicited for as long as the input pulse was present, indicating that they are both input-output devices that convert a neuronal input into a secretion rate of a hormone. These forms of regulation enable immediate response to a specific signal that is introduced into the system (in the form of hormone secretion). In this respect it is not surprising that included in these two classes are fight or flight hormones (adrenaline, noradrenaline), milk (oxytocin) production in response to a stimulus induced by suckling, and melatonin which regulates the circadian rhythm in response to darkness (for additional examples and references see Table 1).

**Table 1:**
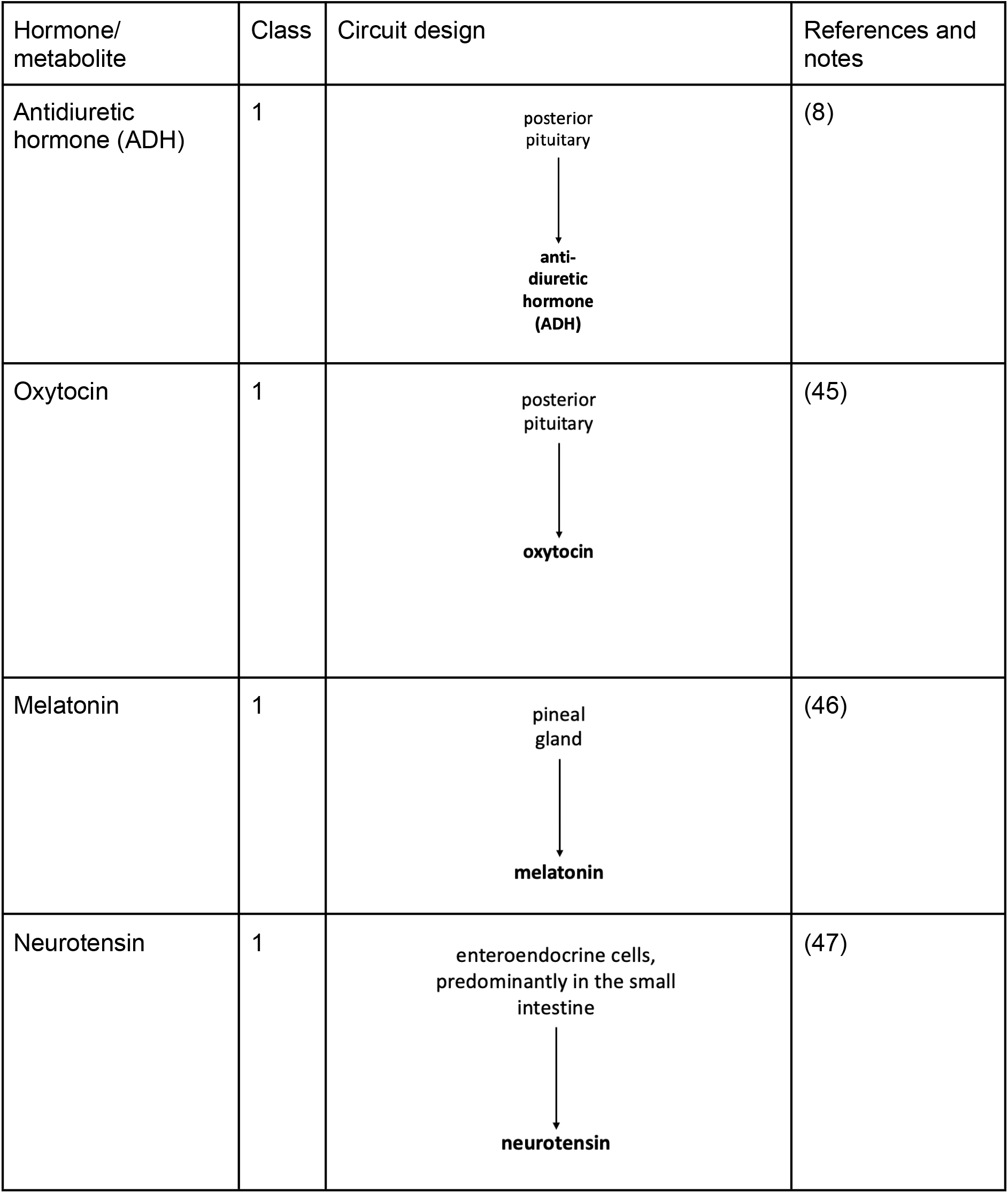
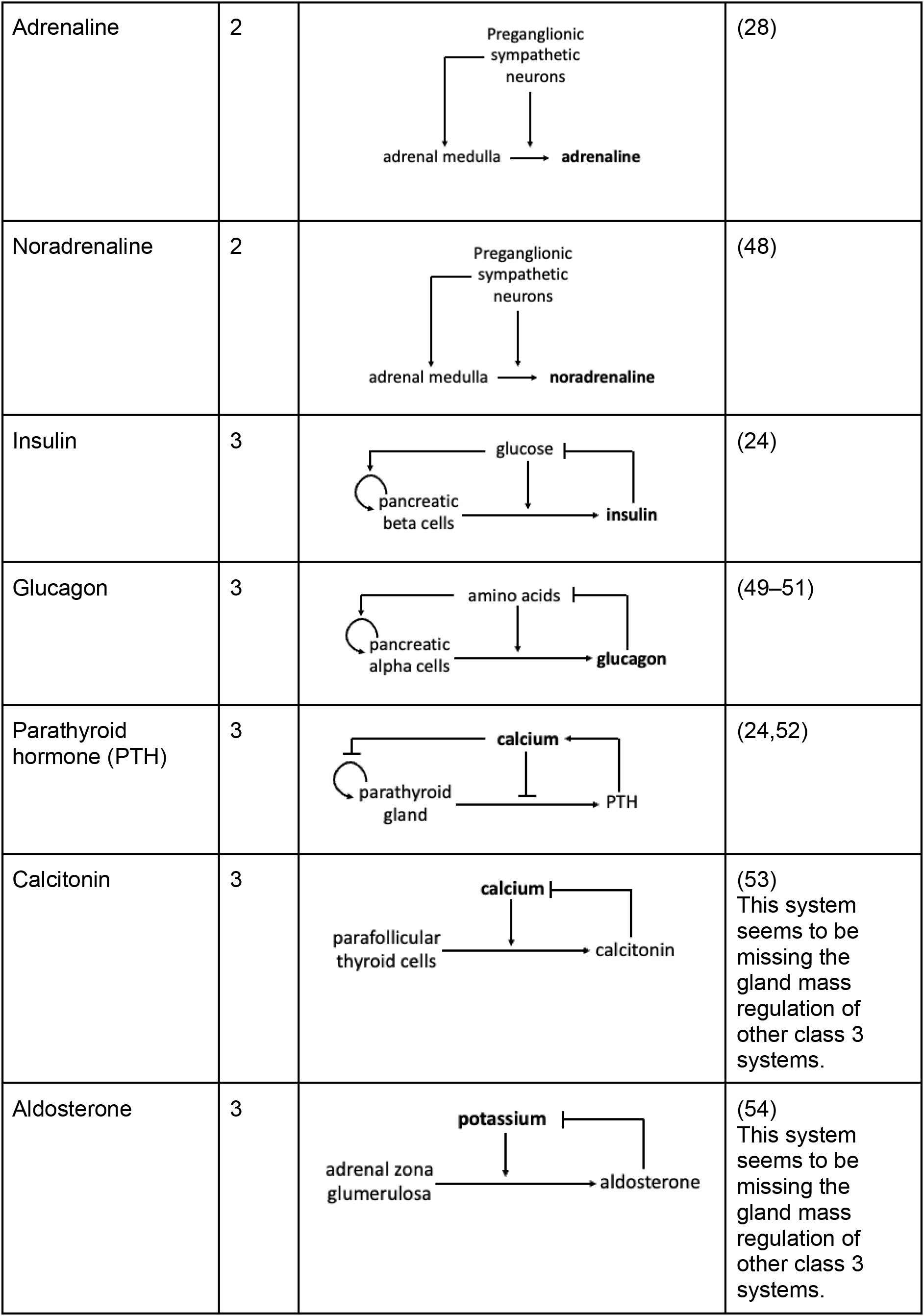
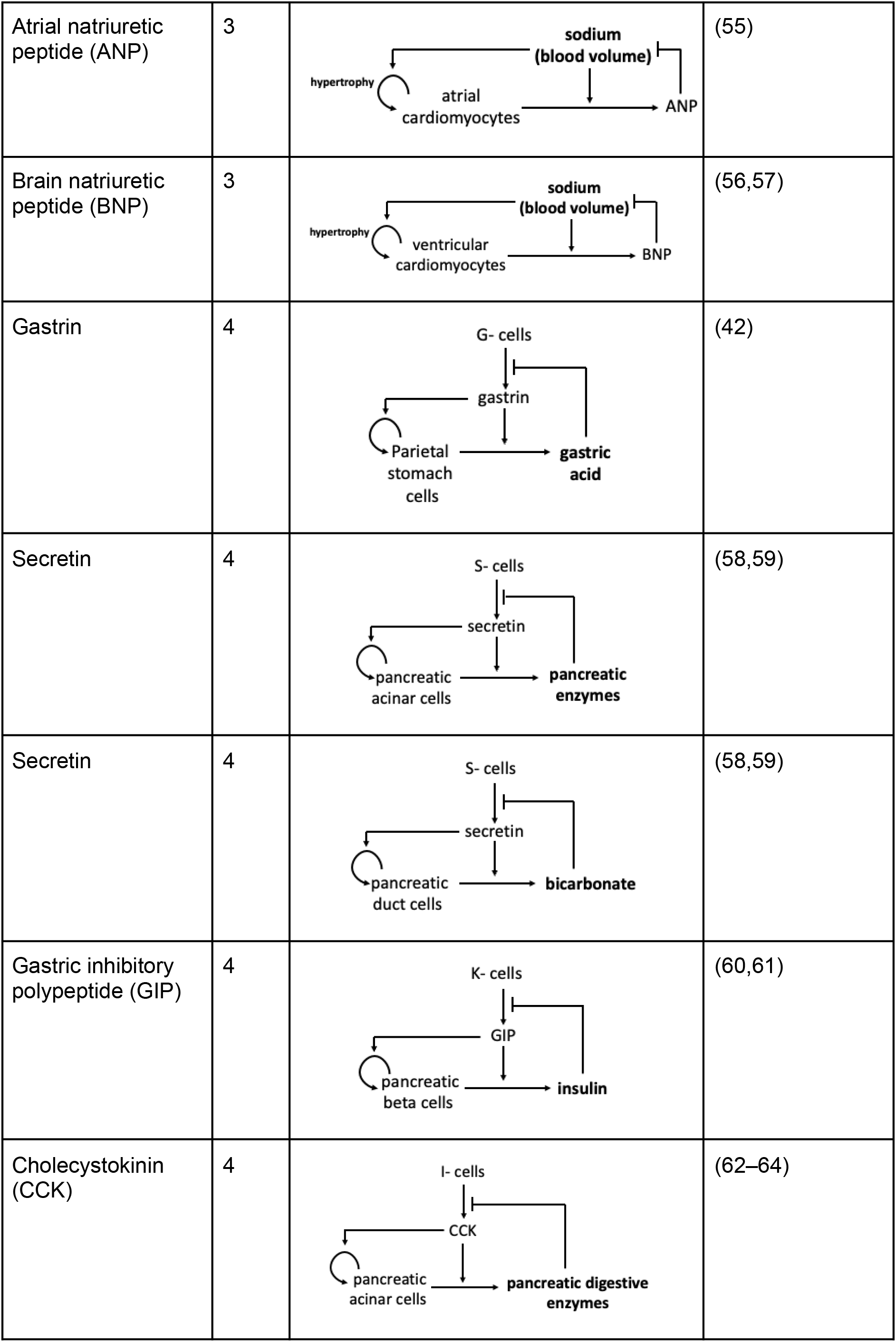
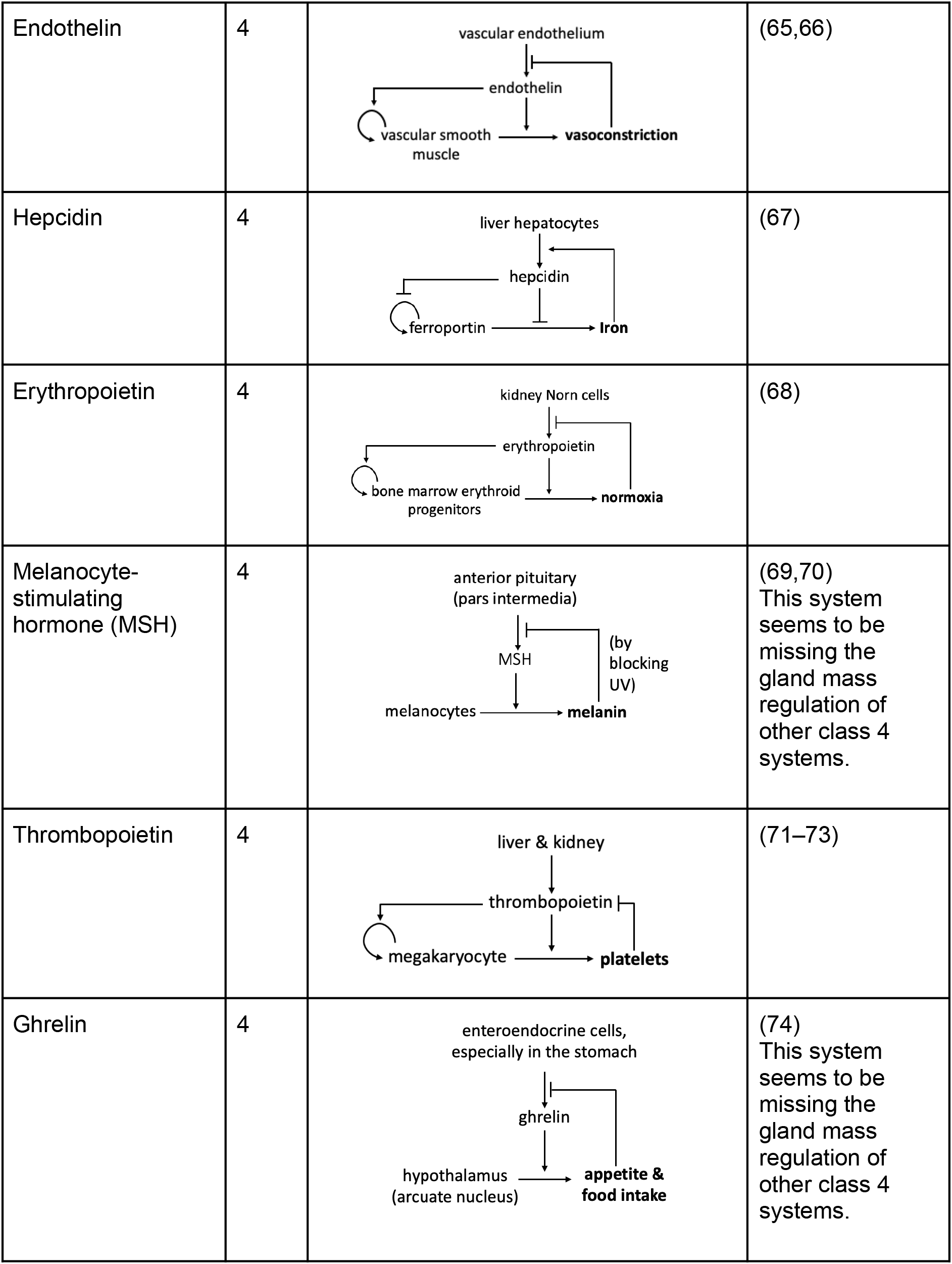
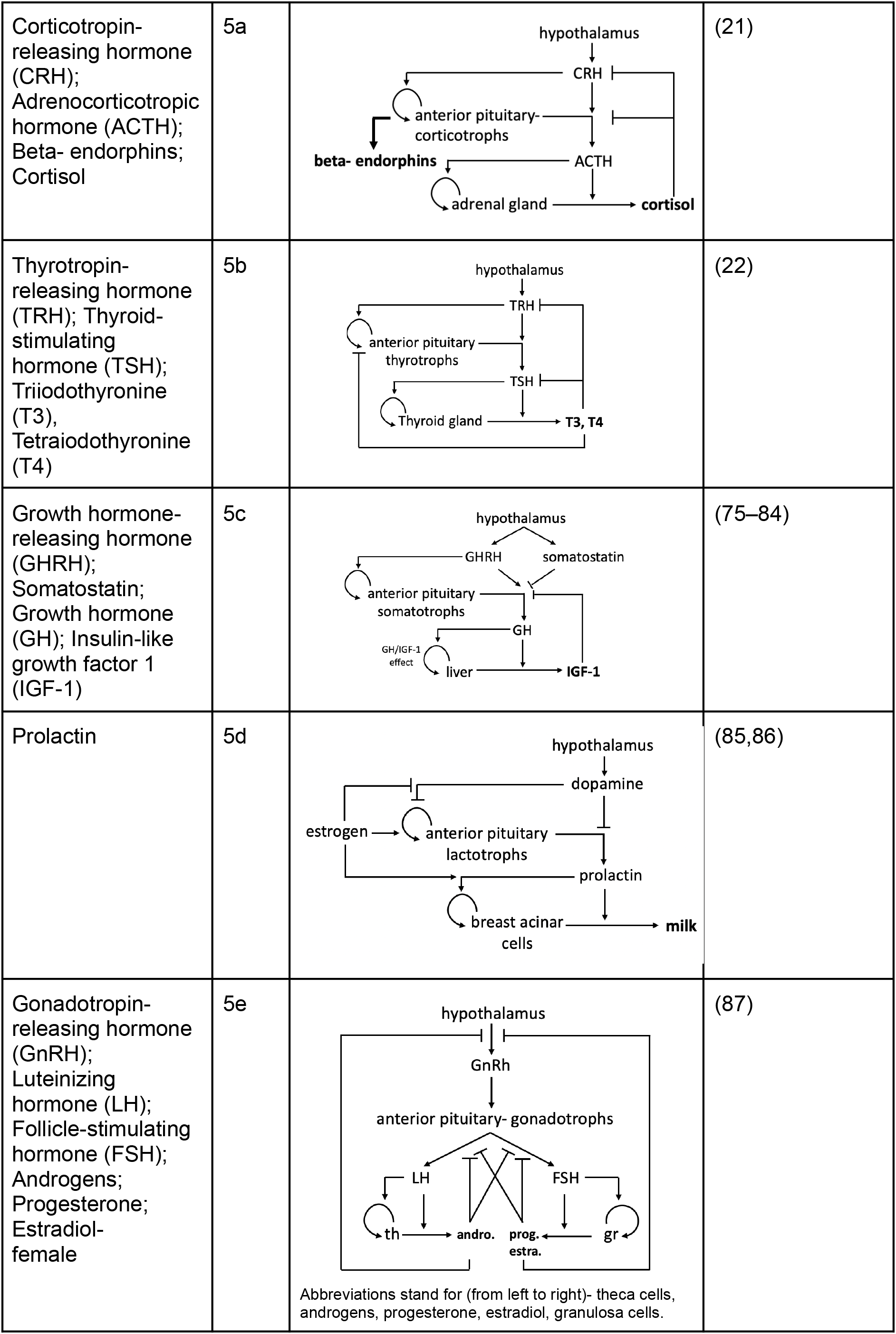
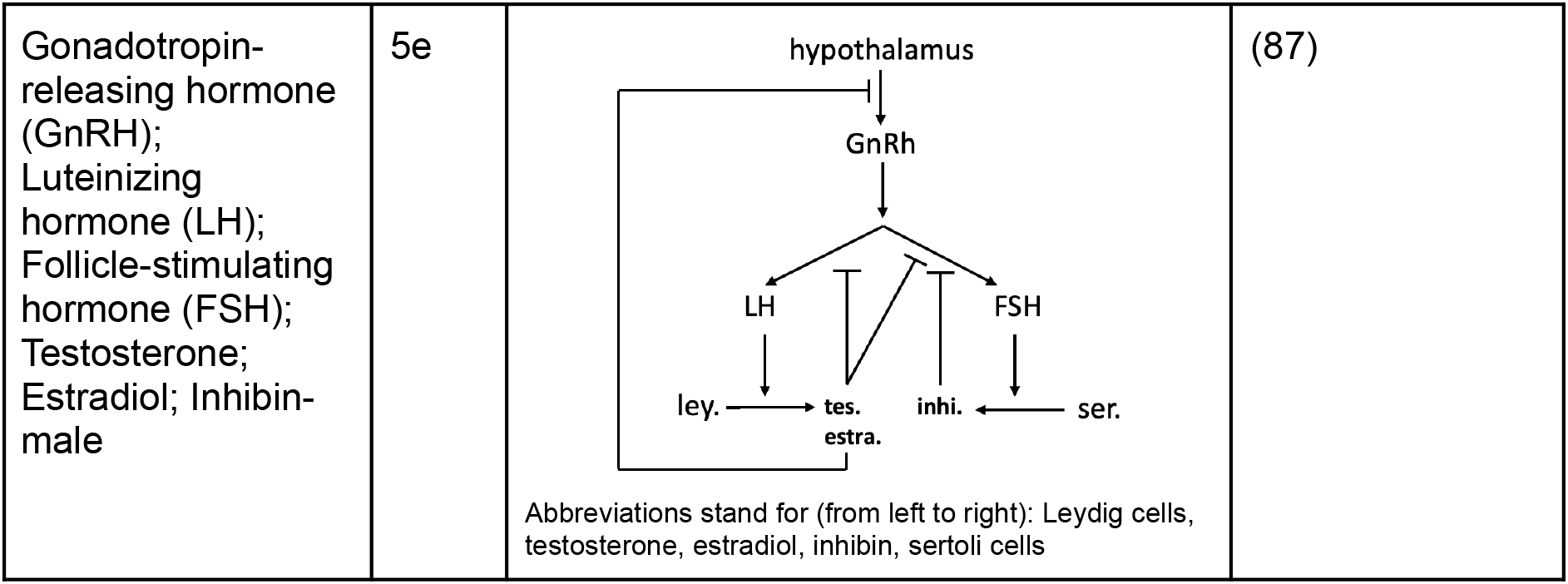
Examples for each endocrine circuit class. The main physiological output of each circuit is in bold.

### Class 3 circuits enable robust homeostasis of metabolite levels

In class 3 circuits, endocrine cells secrete a hormone in response to a metabolic signal (Figure 1c). The hormone acts on the fast timescale of minutes to hours to restore the metabolite to a homeostatic level. The metabolite also regulates endocrine cell growth (hyperplasia or hypertrophy) on the timescale of weeks to months. Examples are beta cells which secrete insulin under control of blood glucose. Glucose is a beta-cell growth signal, acting primarily via hypertrophy in humans after childhood (30,31) and via hyperplasia in rodents(32). Another example is the parathyroid chief cells which secrete parathyroid hormone (PTH) under control of blood-free calcium ions. Free calcium ions also act to regulate parathyroid cell proliferation and survival (33).

Whereas class 1 and 2 circuits provide differing output levels according to inputs, simulations of class 3 circuits showed an ability to achieve a fixed target concentration of a metabolite, as seen in Figure 1c. Class 3 circuits control metabolites with well-defined set points such as 5mM glucose or 1mM calcium ions(34,35). Deviations from these concentrations can be lethal(36,37).

Class 3 circuits are able to act robustly, in the sense that the specific homeostatic concentration of the metabolite is achieved in the face of wide variation in the physiological parameters of the circuit. This is seen in Figure 2a-c, which shows the effect of changing each of the parameters in the circuit. For example, various levels of hormone-sensitivity parameter *s* (such as insulin resistance) lead to the same steady-state metabolite level. Robust homeostasis is achieved by a combination of fast responses and slow gland mass changes(13). The cells respond within minutes to changes in metabolite, such as postprandial insulin secretion. They also respond within weeks by changing their gland functional mass to compensate for physiological changes (Figure 2a-c). As long as the metabolite is away from its set point the cell mass grows or shrinks until the set point is achieved.

**Figure 2:**
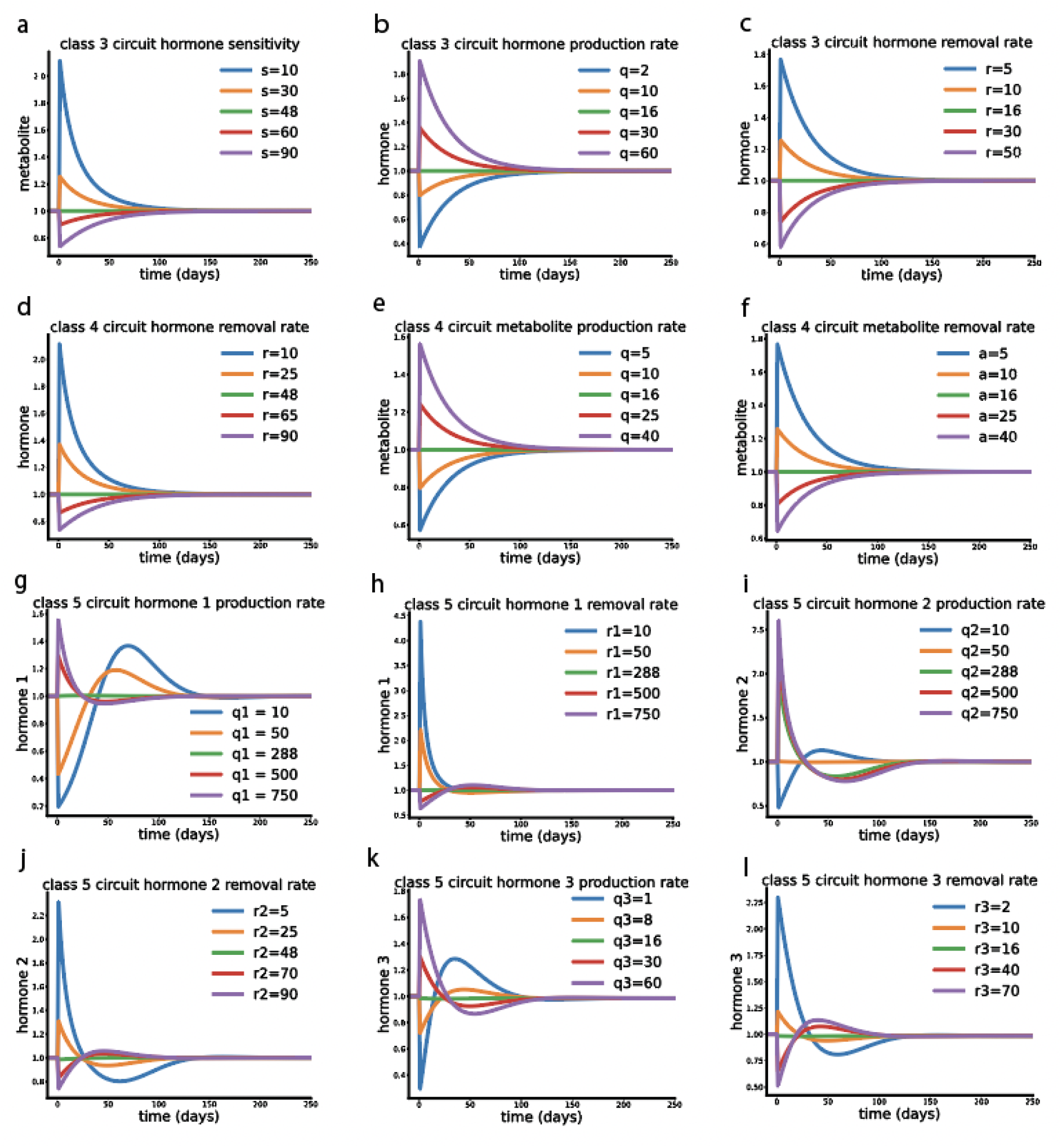
Circuits respond to changes in parameters but their homeostatic variables return to a steady state level that is independent of parameters. Simulations began at steady state with nominal parameter values, and then at *t* = 0 one parameter was changed as indicated, causing a rapid change followed by recovery to the new steady state level, which in all cases is identical to the pre-change level. Green line is no change from the nominal parameter. **(a-c)** Class 3 circuit metabolite and hormone dynamics upon changes in: (a) hormone sensitivity, *s*. (b) hormone production rate, *q*. (c) hormone removal rate, *r*. **(d-f)** Class 4 hormone or metabolite dynamics upon changes in: (d) hormone removal rate, *r*. (e) metabolite production rate, *q*. (f) metabolite removal rate, *a*. **(g-l)** Class 5 circuit dynamics of the three hormones: (g) hormone 1 dynamics with different values of hormone production rate, *q*_1_. (h) hormone 1 dynamics with different values of hormone removal rate, *r*_1_. (i) hormone 2 dynamics with different values of hormone production rate, *q*_2_. (j) hormone 2 dynamics with different values of hormone removal rate, *r*_2_. (k) hormone 3 dynamics with different values of hormone production rate, *q*_3_. (l) hormone 3 dynamics with different values of hormone removal rate, *r*_3_. Parameters not varied in this figure are as in Figure 1. See Box 1 for model equations.

An example is the hypertrophy of beta cells seen in individuals with insulin resistance(38). The change in gland mass can compensate precisely for changes in physiological parameters, as long as the gland is not limited by a maximal size known as a carrying capacity(22,39,40).

Mathematically, the reason for robustness is the integral feedback loop that controls the mass of the secreting cells(13,24) – see the model in Box 1. The equation for the change in gland mass *G* regulated by the metabolite *m* is *dG*/*dt* = *G* (*cm* − *d*), where *c* and *d* are production and removal rates of gland mass. Since *G* is made of cells, it multiplies both production and removal terms (all cells come from cells). This equation ensures that as long as there are secreting cells (*G*≠0), the only possible steady-state of the metabolite is *m*_*st*_ = *d*/*c*. This steady-state solution for the metabolite *m*_*st*_ does not depend on any of the other model parameters, including the metabolite input *u*, its sensitivity *s*, the hormone production rate *q* or the hormone removal rate *r* (Figure 2a-c).

Class 3 systems include, in addition to glucose and calcium control, hormones that regulate electrolytes in the blood that must be kept within a tight range such as potassium and sodium. For more details and examples see Table 1.

### Class 4 circuits enable allostatic control with adjustable set points

Class 4 circuits (Figure 1d) describe secretory cells whose input signal is a hormone, rather than a metabolite as in class 3. These cells secrete a metabolite under control of the input hormone. The input hormone is also a growth factor for the cells on the slow timescale (weeks-months). The input hormone is itself secreted by another cell type. For example, stomach parietal cells secrete acid under control of the input hormone gastrin, which is secreted by other cells in the digestive tract called G-cells that sense neuronal and nutritional signals associated with meals(41). Gastrin is also a growth factor for the gastric parietal cells(42).

Simulations show that this circuit can provide allostasis – the output metabolite has a steady-state set point that can be tuned to physiological needs (Figure 1d). This tuning is achieved by changing the secretion rate of the input hormone and other parameters.

Class 4 circuits lock the input hormone to a homeostatic value on the slow timescale. In a certain sense, the class 4 circuit is the “inverse” of class 3. In class 3, a metabolite achieves homeostasis by allostasis of a hormone. In class 4, a metabolite achieves allostasis by homeostasis of a hormone.

The difference in behavior between class 3 and 4 is due to the position of the hormone in the circuit (see also equations in Box 1). The input signal in class 3 is a metabolite, and in class 4 it is a hormone. When a signal controls the cell growth rate, it participates in an integral feedback loop that locks the input signal to a fixed point on the scale of weeks. In class 3 circuits, metabolites are thus locked to a constant value (e.g., 5mM in the case of glucose(34)). In class 4 circuits, the input hormone is locked, but the output, such as stomach acid, depends on the gland mass, which can adjust to varying parameters on the slow timescale. This steady-state solutions do not depend on any of the other model parameters, including hormone removal rate *r*, the metabolite production rate *q*, or the metabolite removal rate *a* (Figure 2d-f).

We found that enteroendocrine hormones, such as gastrin and secretin, often have a class 4 circuit design. This makes sense as their corresponding metabolites have adjustable set points according to different digestive needs(41). For additional examples see Table 1.

#### Box 1

**The five classes of hormone circuits.**

Circuit diagrams and model equations for the hormone regulatory motifs. *h* = hormone concentration, *m* = metabolite concentration, *G* = endocrine gland mass, *u* = input signal, *r* = hormone removal rate, *q* = hormone/metabolite production rate, *s* = hormone sensitivity, *a* = metabolite removal rate, *c* = cell proliferation/growth rate, *d* = cell death or removal rate.

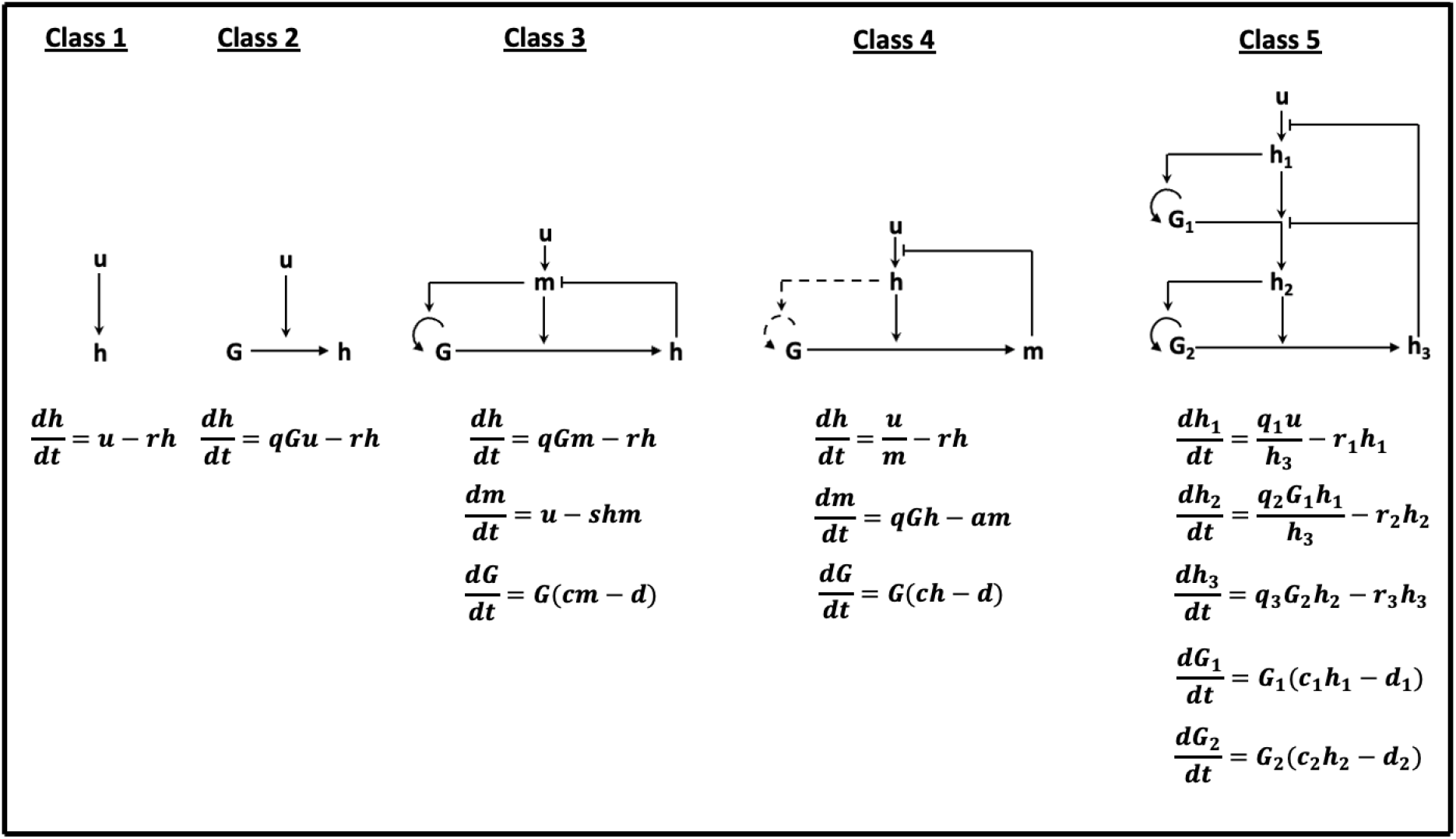

### Class 5 circuits have several functional advantages

Class 5 circuits (Figure 1e) are the most complex. They involve three glands, a top hypothalamic gland and two downstream glands – a pituitary cell type and an effector gland (effector glands can come in pairs as in the right and left adrenals, which we consider as a single gland for our purposes). The pituitary and effector glands can change their functional mass according to the upstream and downstream hormones that act as growth factors. The mass changes are on the slow timescale of weeks-months. This circuit resembles two instances of a class 4 circuit placed on top of each other in series.

The size of the glands shows a hierarchy where the top neuronal gland is the smallest, the middle (pituitary) gland cell type is of intermediate size and the effector gland is the largest. This is due to the principle that gland mass is proportional to its target mass (43). Class 5 circuits are found in the hypothalamic-pituitary axes. These axes control major functions in vertebrates. The hypothalamic-pituitary-adrenal (HPA) axis controls stress response via the hormone cortisol(19). The thyroid axis controls metabolic rate via thyroid hormones(44). Similarly, the sex hormone and growth hormone pathways share a class 5 design.

There are minor variations in design between these axes, which can be considered as several subtypes of class 5 circuit. In the thyroid axis, the growth factor for the pituitary is not its upstream hormone but instead the effector hormone (see Table 1 for details and references) (22).

Simulations of the generic class 5 circuit showed complex dynamical properties. The end product of the circuit (the effector hormone 3 in Figure 1e) shows allostasis and overshoots. The overshoots speed up the response to changing levels of input hypothalamic signal. The upstream hormones 1 and 2 both show homeostasis. The position of hormone 1 and hormone 2 in circuit 5 is analogous to the position of the hormone in circuit 4 and the metabolite in circuit 3, and as the simulations showed, their secretion rate is maintained under homeostatic regulation.

The class 5 circuit also shows robustness properties with respect to changes in its parameters (Figure 2g-l). The levels of *h*_1_ and *h*_2_ hormones are independent of most of the model parameters. The effector hormone *h*_3_ depends linearly on the input *u*, as appropriate for axes that transduce a brain input to a proportional physiological output(13,21).

Due to the complexity of class 5 circuits and centrality in hormone regulation, we next consider their functions in more detail in the remainder of this study.

### The pituitary serves as an endocrine amplifier

Next, we explore the structure-function relationship of class 5 circuits. One question is what advantage this design, with a pituitary gland in the middle, might have compared to simpler circuits that lack a middle gland. Many of the hormones that are secreted from the pituitary, such as ACTH, TSH, LH and FSH, do not have a major physiological role except to serve the next step in the circuit, inducing the effector gland to secrete the effector hormone (thyroid hormones in the case of TSH, cortisol in the case of ACTH and sex hormones in the case of LH and FSH). This can be seen from the expression pattern of the receptors for these hormones in the human body (88–93). In the next few sections, we examine the role of the pituitary and offer several functional advantages that are absent from designs that lack a middle gland.

We propose that one function of the pituitary is to act as an amplifier of the hypothalamic hormones. We showed in previous work that a single endocrine cell serves about 2000 target cells on average, regardless of the hormone in question (43). In the Hypothalamic-Pituitary (HP) class 5 circuits, a small brain region, the hypothalamus, secretes hormones. If there were no pituitary, the hypothalamus would need to secrete enough hormones for the effector gland, such as the adrenal, thyroid and female and male gonads. These glands in turn serve nearly the entire body, 5 × 10^12^ cells(94), and thus have about 10^9^ cells(43). Without a pituitary, the hypothalamic regions that secrete each hormone would thus need to have a mass of about 10^6^ cells, which is 2 orders of magnitude larger than observed for the neuronal cells secreting CRH, TRH and GnRH (we could not find estimates for the number of GHRH secreting neurons) (43,95,96). It may be implausible to host such a large number of cells in the hypothalamus. The pituitary, which lies external to the skull, can more easily accommodate a large number of endocrine cells. It may thus have an amplification role, allowing 10^4^ hypothalamic cells to produce enough hormone for the 10^7^ pituitary cells(11,43,97), which then provides enough hormones for the 10^9^-10^10^ cell effector gland. For scale, 10^9^ cells typically weigh 1 gram (98) (Bionumber ID 111609).

### The pituitary can compensate for toxic adenomas up to a threshold

Beyond this amplification role, we asked whether the pituitary also has dynamical functions. Another way to pose this question is to ask why the pituitary is regulated to change its functional mass in class 5 circuits, rather than having a constant mass.

We begin by noting that changes in the pituitary mass can buffer physiological and pathological variations. An example has been described previously in the context of thyroid disease (22). We add to this previous work an exploration of how class 5 circuits might compensate for tumors that hypersecrete hormones.

As a concrete example we consider the HPA axis in which tumors arise quite frequently, in up to a few percent of the population (often called incidentalomas) (99). They occur both in the pituitary cells that secrete ACTH and in the adrenal cells that secrete cortisol (100,101). When the tumors occur, the corresponding non-tumorous gland mass shrinks(105,106). The tumors often have no physiological consequence – the levels of the target hormone cortisol are normal (102,103). When tumors that secrete hormones exceed a threshold size, they dysregulate the hormone levels, causing overt hypercortisolism called Cushing’s syndrome. The tumors thus transition from a subclinical to a clinical disease.

We asked what sets the threshold between subclinical and clinical disease. We also asked whether there are qualitative differences in the dynamics between tumors in the pituitary and tumors in the adrenal that have the same net effect of increasing cortisol levels. To address this, we used the mathematical model for the HPA axis from Karin et al.(21), which represents a special case of the class 5 equations in Box 1 (Methods). We updated the equations to include the growth of the tumor and its effect on cortisol production. We modeled the growth of the tumor *C* with an additional equation *dC*/*dt* = *rC*(1 − *C*/*K*) describing logistic growth at rate *r* with carrying capacity *K*. We make the simplifying assumption that the carrying capacity depends only on constant features like anatomy and not on dynamic variables like cortisol. We modeled cortisol production γ per unit tumor mass by adding a term γ*C* to the equation describing cortisol dynamics *dh*_3_/*dt*. We analytically solved the updated HPA class 5 mathematical model at steady state (Methods).

We begin with an adrenal tumor that secretes cortisol (Figure 3a). We assumed, as commonly observed, that the tumor secretion rate is not regulated by ACTH(104). We find that, as long as the normalized tumor secretion rate β is below a critical threshold, the non-tumorous adrenal mass shrinks to *precisely* compensate for the extra hormone secreted by the tumor (Figure 3a-d).

**Figure 3.**
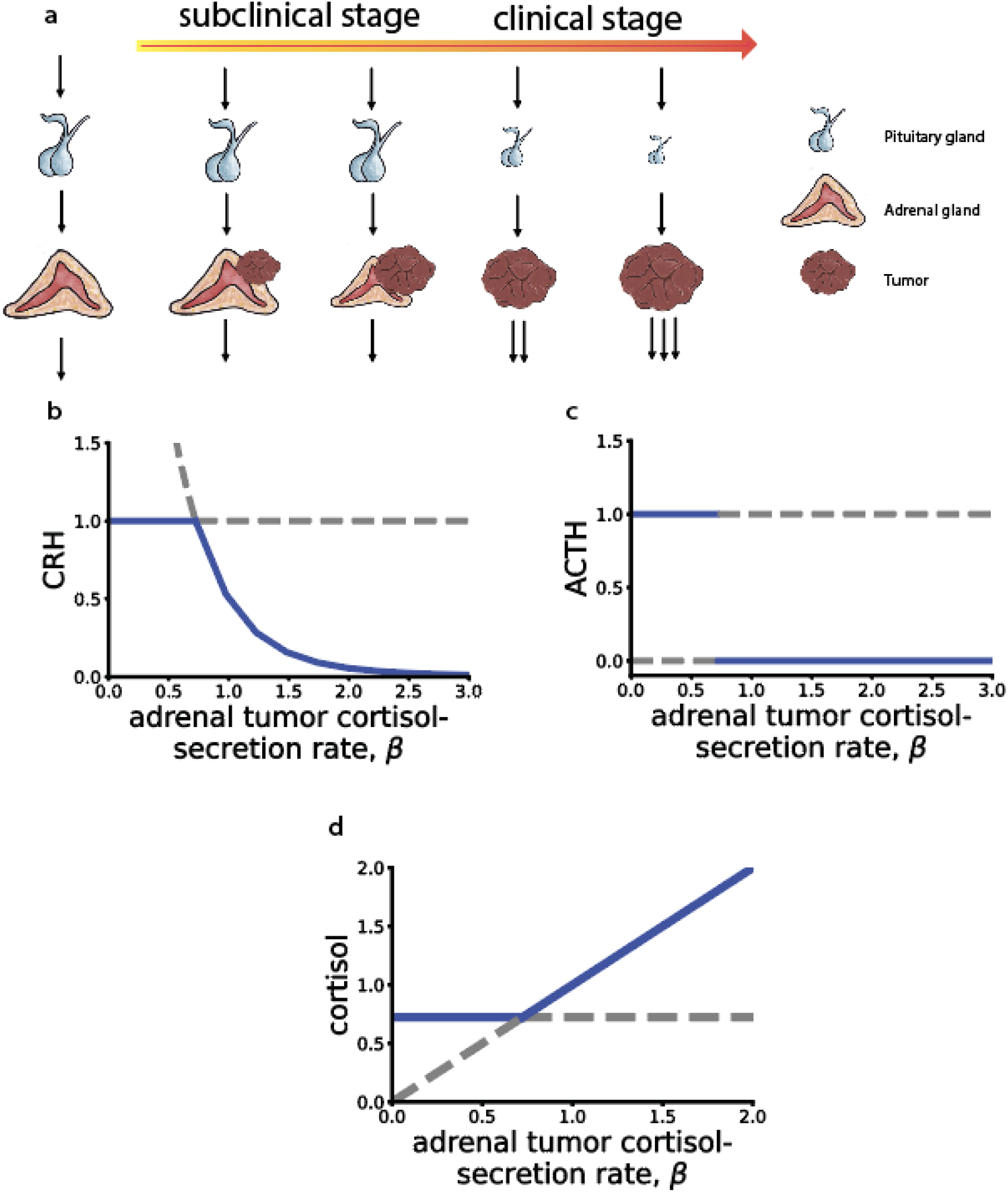
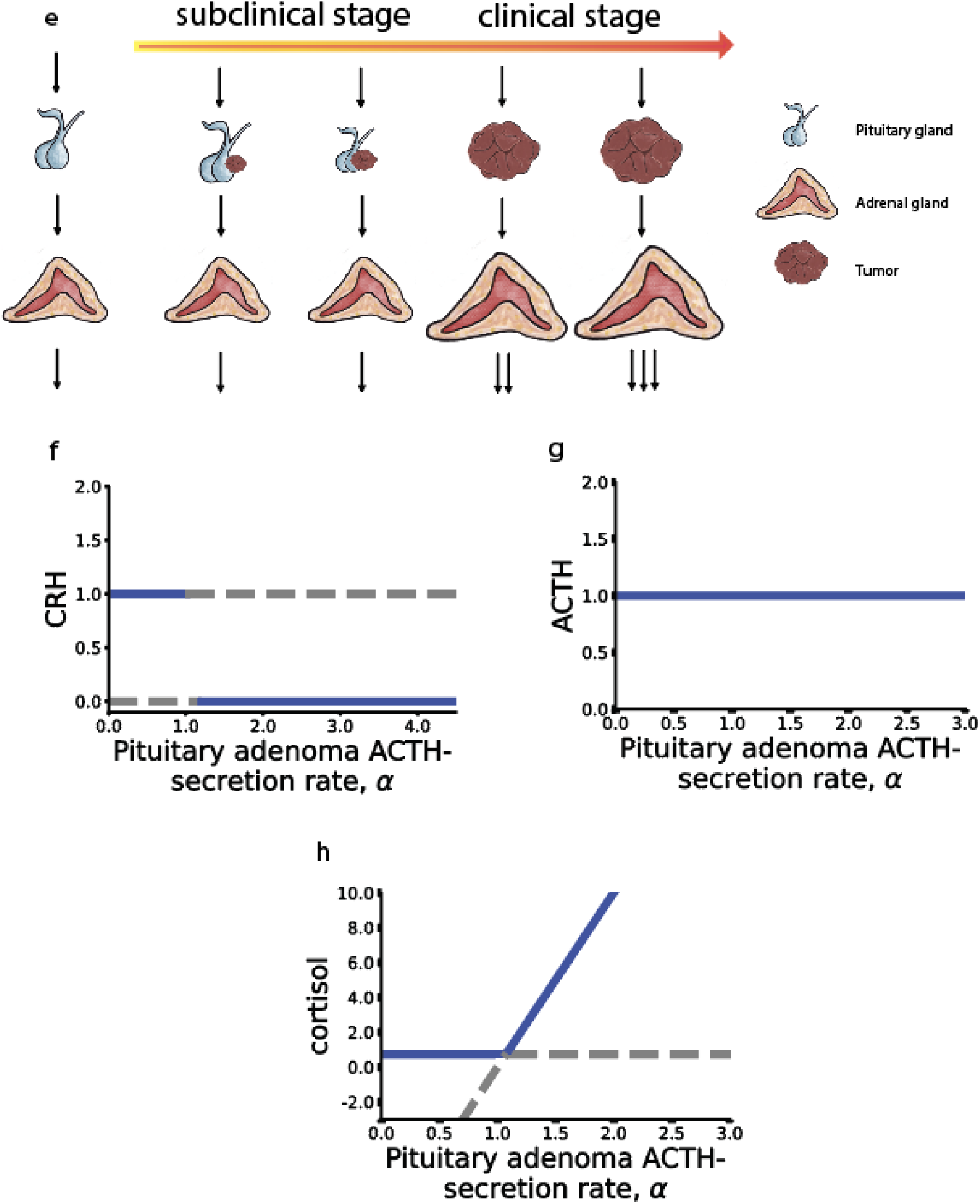
Transition between subclinical and clinical Cushing’s syndrome cases for pituitary and adrenal tumors. **(a)** Schematic showing how an adrenal cortisol-secreting tumor causes the native (non neoplastic) adrenal cortex cortisol-secreting mass to shrink. Clinical disease appears in cases in which the adrenal shrinks to zero. **(b)** CRH **(c)** ACTH and **(d)** cortisol as a function of adrenal tumor cortisol secretion rate β. **(e)** Schematic showing how a pituitary ACTH-secreting tumor causes the native pituitary corticotroph mass to shrink. Clinical disease appears in cases in which the pituitary shrinks to zero. **(f)** CRH **(g)** ACTH and **(h)** cortisol as a function of pituitary tumor ACTH secretion rate α. We simulated using *q*_1_ = *r*_1_,= 288 *day*^−1^, *q*_2_ = *r*_2_ = 48 *day*^−1^, *q*_3_ = *r*_3_ = 16 *day*^−1^, 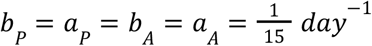, *r* = 1 *day*^−1^, *K*_*gr*_ = 4, *n* = 3, *K* = 1, *u* = 1, and ζ = 0. 1. See Methods for model Equations 7 – 12 for the simulations in (b-d) and Equations 31 – 36 for the simulations in (f-h).

However, at a critical secretion rate, the native (non-tumorous) adrenal functional mass shrinks to zero at steady state (a transcritical bifurcation). This critical rate occurs when the tumor produces exactly the same amount of cortisol as the native pre-tumor adrenal glands. Thereafter, as the tumor secretion grows, cortisol levels exceed normal levels (Figure 3a-d) and clinical symptoms of hypercortisolism occur, called Cushing’s syndrome. Since tumor growth typically evolves over years, whereas glands adjust over months, we assume quasi-steady state and simulate the circuit with a constant value of total cortisol secretion β. This represents a very slow change in β or a non-temporal distinction between individuals with subclinical and clinical tumors.

We compared this adrenal tumor to a different form of Cushing’s pathology in which a pituitary tumor secretes ACTH. To address this, we used the mathematical model from Karin et al.(21) and updated the equations to include the growth of the tumor and its effect on ACTH production. We modeled the growth of the tumor *M* with an additional equation *dM*/*dt* = *rM*(1 − *M*/*K*) describing logistic growth at rate *r* with carrying capacity *K*. We again make the simplifying assumption that the carrying capacity depends only on constant features like anatomy and not on dynamic variables like cortisol. We modeled ACTH production η per unit tumor mass that is suppressed by cortisol by adding a term to the equation describing ACTH dynamics *dh*_2_ /*dt*. We find that at a secretion rate α below a critical value, the native (non-tumor) pituitary corticotroph mass shrinks to compensate and maintain a normal level of ACTH and cortisol due to the negative feedback of cortisol secreted from the adrenal, as known from the literature(1,107) (Figure 3e-h). However, beyond a critical tumor secretion rate, the non-tumor pituitary corticotroph functional mass shrinks to zero at steady state. The critical secretion rate occurs when the tumor secretes the same amount of ACTH as the pre-tumor pituitary corticotrophs. Thereafter, higher tumor secretion causes elevated levels of cortisol, resulting in Cushing’s disease (Figure 3e-h). As above we assume a quasi-steady state of very slow tumor growth compared to gland mass changes and hence constant α.

In both ACTH– and cortisol-secreting tumors, the system undergoes a transition which, in the language of dynamical systems, is a transcritical bifurcation(108). Beyond the transition, the compensating gland functional mass (the non-tumor part of the gland) drops to zero. We conclude that changes in gland mass can compensate for toxic adenomas until they reach a critical mass.

### The pituitary can speed responses on the scale of weeks compared to simpler circuits

We also asked whether the pituitary can provide dynamical benefits to class 5 circuits as compared to simpler class 1-4 circuits in non-pathological situations. For this purpose we studied the HPA model, using the equations from Karin et al. from above(21) (Methods). We tested this model with a prolonged stress input that rises and remains high for months and then falls, to explore the onset and recovery from such prolonged stress(21). The reason we chose to analyze the dynamics of the HPA axis during stress is because stress and high cortisol secretion are expected to occur during health (stressful work periods, migration, lack of proper sleep due to childbearing, etc.) and therefore studying the dynamics of the system before, during and after such prolonged stress periods can reveal dynamical properties of its function.

We compared the class 5 HPA circuit with two hypothetical simpler designs for cortisol control. One has a pituitary but the pituitary cannot change its mass. We thus set *dP*/*dt* = 0 and *P* = *P*_0_. The second alternative circuit is a class 4 circuit without a pituitary. To model this we dropped the *dh*_2_ /*dt* and *dP*/*dt* equations and replaced *h*_2_ with *h*_1_. Here the adrenal is directly activated by a hormone from the hypothalamus. To allow a ‘mathematically controlled comparison’(13,109,110) we set all hormone half lives and production rates to be equal between the circuits.

We simulated these three circuits and their responses to prolonged (500 days) stress input (Figure 4, Methods). In the class 5 HPA circuit (Figure 4a,d,e), the onset of stress causes a rapid response on the scale of hours, and then a slower increase on the scale of weeks as gland masses change(21). Similarly, at the end of the stress input pulse, there is a rapid reduction on the scale of hours, followed by a slower adjustment due to gland mass changes on the scale of weeks.

**Figure 4.**
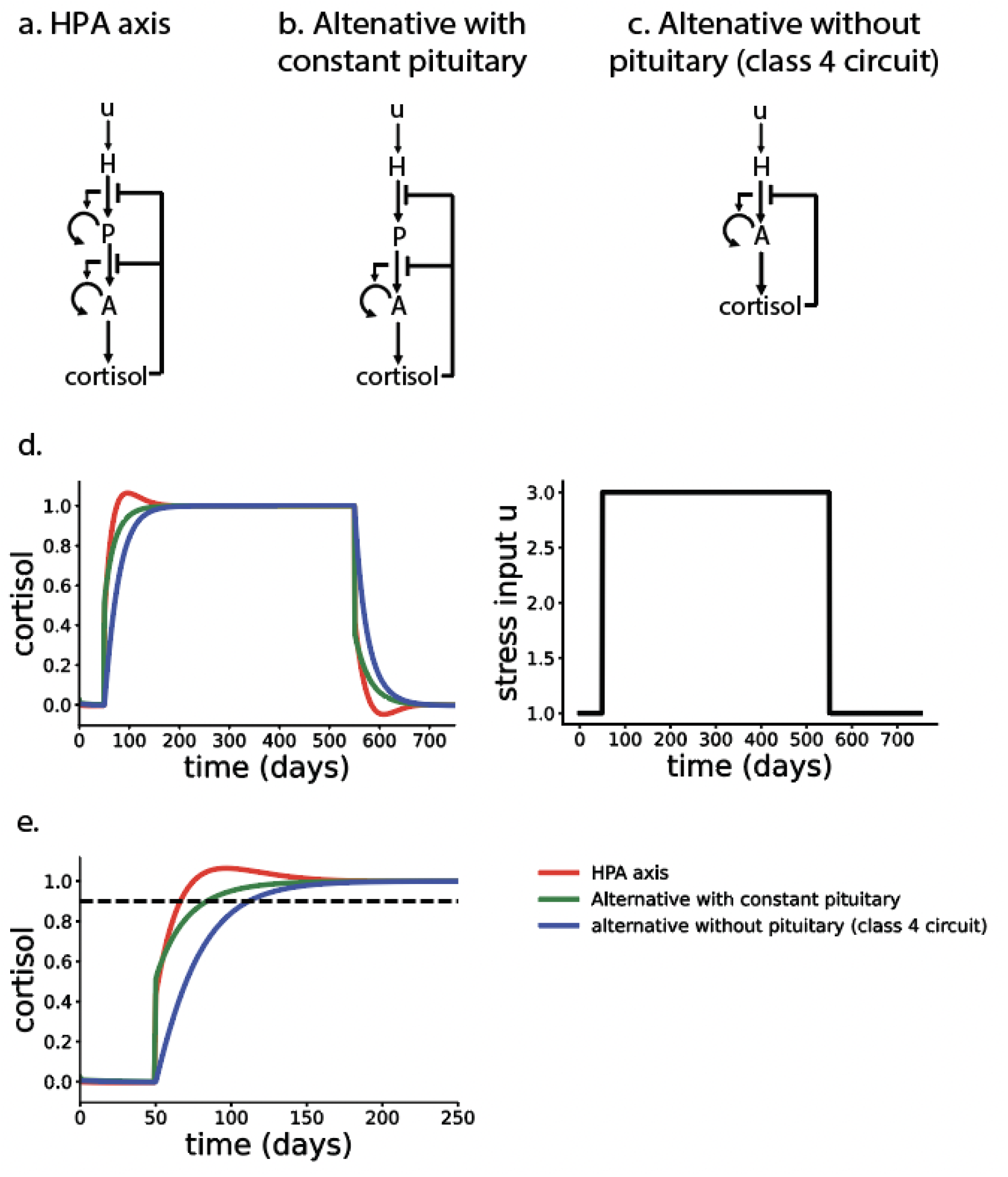
Comparison of alternative HPA designs indicates that the natural design overshoots and has the fastest rise time. **(a)** natural class 5 design of the HPA axis in which the pituitary functional mass is regulated, **(b)** alternative design with a pituitary that has a constant functional mass **(c)** alternative design without a pituitary (no middle gland). **(d)** Dynamics of cortisol in response to a prolonged stress input which rises to 3 times the normal input and drops back 500 days later. **(e)** Zoom-in on the rise phase shows that class 5 circuit reaches 90% (dashed black line) of steady-state response fastest. We simulated with parameters *q*_1_ = *r*_1_ = 288 *day*^−1^, *q*_2_ = *r*_2_ = 48 *day*^−1^, *q*_3_ = *r*_3_ = 16 *day*^−1^, 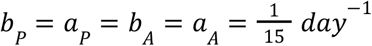 *K*_*gr*_ = 4, *n* = 3, *u* = 1. See Methods for the model equations used in the simulations.

The alternative class 5 circuit with constant-mass pituitary (Figure 4b,d,e) has a 35% slower rise time, defined as the time to first reach 90% of the hormone steady state value. The alternative class 4 circuit without a pituitary (Figure 4c,d,e) has the slowest rise time (75% slower). The same is found upon recovery from the stress pulse – the class 5 circuit reaches within 10% of the baseline faster than the other circuits. The class 5 circuit, unlike the alternative ones, also shows an overshoot due to the adjustment of pituitary mass absent from the other two simpler circuits. We conclude that the pituitary with changing mass can provide a speedup on the scale of weeks when stress conditions change.

## Discussion

We analyzed literature data on 43 human hormones and modeled them mathematically. We found that endocrine circuits can be organized into 5 classes according to recurring regulatory circuit motifs. The design of each class of circuits serves a specific function such as acute quantitative response, endocrine amplification, strict homeostasis or adjustable set points, as we demonstrate using simulations and analytical solutions. Regulation has two timescales, a fast secretion timescale of minutes to hours, and a slow adjustment of gland functional mass on the scale of weeks, that allows robustness to physiological variations and the possibility of different set points.

The five classes of circuits defined here have characteristic functional roles. Class 1 and 2 circuits can serve as fast input-output devices. In class 1, specialized neurons directly produce hormones, whereas in class 2, neurons activate hormone-secreting glands. We speculate that class 2 may allow for a more nuanced or quantitative response compared to class 1, which may primarily reflect all-or-none output. Class 3 circuits can lock a metabolite to a tight range around a specific concentration. An example is control of glucose tightly around 5mM and control of blood free-calcium ions around 1mM(34,35). Class 4 circuits lock the input hormone level and can offer allostatic control of their output metabolite. Examples are intestinal hormones secretin and gastrin that control bicarbonate secretion and stomach acid secretion respectively(42,58,59). Finally, class 5 circuits are the most complex. They can amplify hormones to levels that serve the entire body, speed responses and compensate for small toxic tumors and for variations in physiological parameters such as hormone production rates.

The mass of endocrine glands can expand by hypertrophy or hyperplasia, depending on the gland and on age(22,24). It can also grow, in principle, by differentiation of progenitor cells(111). One basic function of such mass growth occurs when high levels of hormones are needed for long times. The circuits of classes 3-5 can sense this need and signal the endocrine cells to grow in total mass and thus in their hormone secretion potential. A well-known example is endemic goiter in which the thyroid expands when iodine, essential for thyroid hormone production, is very low(112). The thyroid mass can grow by a factor of ten or more. The thyroid also grows in pregnancy to meet the needs of the fetus(113). Similarly, the adrenal cortex grows in people under chronic stress or depression(114,115).

Another consequence of the gland mass regulation by these circuits is that they offer a solution to the problem of organ size control. Since cells expand exponentially, their growth and removal rates need to be precisely matched to avoid excess mass growth or shrinkage(40). The circuits ensure that the gland mass production and removal rates balance precisely when the signal reaches a functional level(24). Thus, the same circuits solve two problems: expansion of gland mass when more hormone is needed, and organ size control.

One interesting question in class 5 circuits is the purpose of a middle gland – the pituitary in HP axes. Why would an alternative design without a middle gland, like class 4 circuits, not be chosen by natural selection instead? We propose several answers. First, the middle gland acts as an amplifier. In the hypothalamic-pituitary (HP) axes, a small brain region in the hypothalamus secretes a hormone, and the effector gland needs to serve the entire body, and is thus on the order of 10^9^ cells as found in the mass law of (43) where each endocrine cell serves about 2000 target cells. In order to supply such a large effector gland, it is useful to have a middle gland (eg 10^7^ − 10^8^ for each secretory cell type) so that the hypothalamic region can be small. A tiny amount of hypothalamic hormone thus regulates a sub-pea-sized pituitary, which regulates a large effector gland. In the HP axes, there is about a thousand-fold ratio between the top and middle gland, and a similar ratio between the middle and effector gland.

A second functional advantage of the middle gland is speedup of responses to inputs that change on the scale of many weeks. Alternative designs that lack a middle gland, or with a middle gland that can’t change its mass, have slower response on the scale of weeks. The middle gland in class 5 circuits achieves speedup by causing a mild overshoot or undershoot of several weeks in the hormone dynamics. This overshoot can cause mild dysregulation when entering or exiting prolonged periods of high excitation, as in prolonged stress(21), postpartum(25) or after addiction(26).

Finally, the middle gland can participate in compensation of physiological or pathological perturbations. We demonstrate this by analyzing the effects of hormone-secreting tumors in the HPA axis that cause Cushing’s syndrome. A cortisol-secreting tumor in the adrenal can be fully compensated by reduction of the rest of the adrenal cortisol-secreting mass (caused by a reduction in the adrenal growth factor ACTH due to feedback regulation), provided that the tumor secretion rate is below a threshold value. This avoids clinical consequences of mildly cortisol-secreting tumors, which may account for 15% of tumors (incidentalomas) found in the adrenal(101). These mildly secreting tumors are thus quite common subclinical events given that incidentalomas are found in about 4% of individuals undergoing high-resolution abdominal imaging(116). When the tumor secretion rate crosses a threshold value, however, the effective adrenal mass shrinks to zero, and compensation cannot continue. Thereafter, high cortisol with clinical symptoms can result, a condition known as Cushing’s syndrome. The same phenomenon can also be found for ACTH-secreting tumors in the pituitary, which can be compensated by reduction of pituitary functional mass until the native pituitary shrinks to zero and clinical signs of hypercortisolism occur.

It would be interesting to compare these results in humans with other organisms. The major hormone regulatory circuits tend to be conserved in mammals and vertebrates, with the main differences being switches between the dominant chemical form of the hormone (e.g. cortisol in humans, corticosterone in mice) and sometimes in its biological functions on target tissues. It is thus plausible that the same five circuit classes will occur across vertebrates despite variation in biochemistry. The question of the role of gland mass changes in other species also requires further research. Recent advances signal the availability of large-scale data in other species in the near future, such as a study on the hormone network of a primate, the mouse lemur(117).

The present hormone circuits may help to understand the circuits of the 21 hormones not included in this analysis because their regulation is not yet sufficiently characterized. One may hypothesize that these hormones will also fall into the five classes.

In summary, endocrine regulatory mechanisms fall into 5 classes of recurring circuit motifs with specific dynamical functions. These circuits cause gland masses to grow or shrink to compensate for physiological and pathological changes. It would be interesting to explore whether other design principles can be found to deepen our understanding of systems endocrinology.

## Acknowledgements

We thank members of the Alon lab for discussions and comments on the manuscript. Funding was provided by the European Research Council (ERC) under the European Union’s Horizon 2020 research and innovation program (grant agreement No. 856587). D.S.G. was funded as a member of the Zuckerman Postdoctoral Scholars Program. Y.K.K is supported by the JSMF Postdoctoral Fellowship in Understanding Dynamic and Multi-scale Systems (Award #https://doi.org/10.37717/2020-1428). U.A is the incumbent of the Abisch-Frenkel Professional Chair.

## Methods

### Hormone circuit inference

We searched the literature for all known human hormones, primarily using an endocrinology textbook (1) and a medical database (4), and found 64 hormones. Using the literature, we focused on each hormone to understand its regulatory circuitry and gland mass control. For each of the secreting cell types, we sought the major growth factor signals, and the major signals for hormone secretion. We excluded hormones where insufficient information was found, such that we could not construct the regulatory circuit (excluded hormones are: adiponectin, alpha-melanocyte stimulating hormone (alpha-MSH), amylin, angiotensin, apelin, c-type natriuretic peptide (CNP), calcitonin-gene related peptide, galanin, glucagon-like peptide 1 (GLP-1), glucagon-like peptide 2 (GLP-2), leptin, motilin, neurokinin B (NKB), osteocalcin, parathyroid-hormone related protein (PTHrP), pancreatic polypeptide (PP), peptide histidine-isoleucine (PHI), peptide YY (PYY), pituitary adenylate cyclase activating polypeptide (PACAP), renin, vasoactive intestinal polypeptide (VIP)).

### Mathematical modeling

We used the principles in (40) to model the different systems: We used ordinary differential equations with production and first-order removal equations for each hormone and metabolite. We modeled positive and negative regulatory effects of a signal on these rates by multiplication or division respectively. We used equations for cell mass by having cell production and removal be both proportional to mass (all biomass comes from biomass), with a growth rate regulated by signals. Simulations used python 3.9.12.

### Steady state equations for class 5 circuit with tumor growth

#### Adrenal adenoma

We modeled the tumor using the Karin et al. model for the HPA axis(21) and updated the class 5 equations from Box 1 to include the growth of the tumor and its effects on cortisol secretion, yielding the following equations:

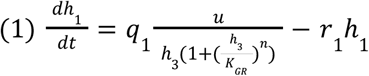

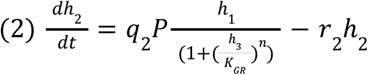

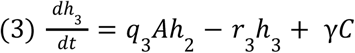

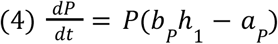

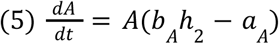

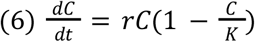

where *h*_1_ is CRH, *h*_2_ is ACTH, and *h*_3_ is cortisol. The parameters *q*_1_, *q*_2_, *q*_3_ represent the corresponding hormone production rates and *r*_1_, *r*_2_, *r*_3_ the corresponding hormone removal rates. The gland parameters *b*_*P*_, *b*_*A*_ are the cell proliferation/growth rates of the pituitary and adrenal, respectively; *a*_*p*_, *a*_*A*_ are the corresponding cell removal/death rates. For the tumor, *r* is the growth rate and γ is the concentration of cortisol secreted per unit mass of tumor *C. K*_*gr*_ is the receptor affinity for cortisol, *n* is the receptor cooperativity for cortisol and *K* is the tumor carrying capacity. Note that the important variable is the total cortisol secretion by the tumor at its final size (beta=gamma K) – one need not assume that gamma or r is constant between individuals or over time, or that each unit mass of the tumor secretes cortisol equally.

We normalized the equations by substituting 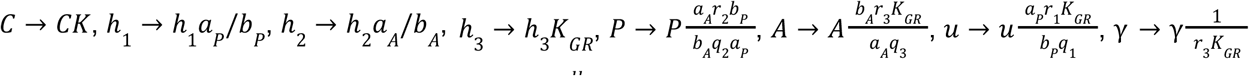 to obtain:

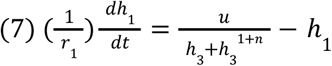

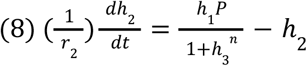

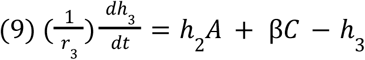

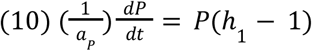

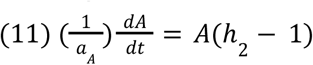

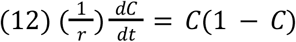

where we define β = γ*K* as the rate of cortisol secretion from the tumor at its carrying capacity, which is the maximal cortisol secretion rate.

At steady state, Equations 7 – 9 have a single solution, whereas Equations 10 – 12 can each be satisfied by two solutions (*P* = 0 or *h*_1_ = 1, *A* = 0 or *h*_2_ = 1, and *C* = 0 or *C* = 1, respectively). We are not interested in situations with *C* = 0, which do not involve the tumor at all.

Steady state solutions in normalized form with β below the bifurcation –– defined implicitly by 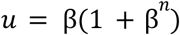 –– are:

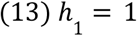

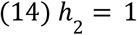

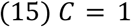

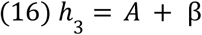

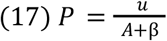

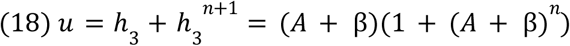

Here the native adrenal functional mass *A* > 0 shrinks to compensate for the tumor, and the tumor reaches its carrying capacity *C* = *K*.

Steady state solutions in normalized form with β above the bifurcation –– defined implicitly by 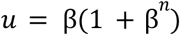 –– are:

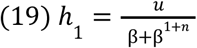

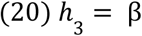

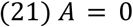

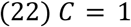

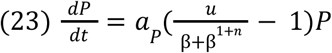

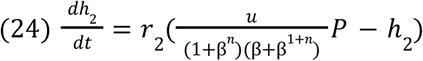

Here native adrenal functional mass = 0, and the tumor reaches its carrying capacity *C* = *K*.

While Equations (19 − 22) are in steady state, we leave Equations (23), (24) as differential Equations to show the time dependence on the external input *u* and the rate of cortisol concentration secreted from the tumor at carrying capacity β. If external input *u* > β + β^1+*n*^, then *P* → ∞ and *h*_2_→∞.If external input *u* < β + β^1+*n*^, then *P* → 0 and *h*_2_→0.

### Pituitary adenoma

We modeled the tumor using Karin et al. model for the HPA axis(21) and updated the equations to include the growth of the tumor and its effects on cortisol secretion, to yield the following equations:

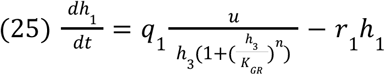

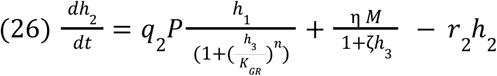

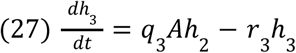

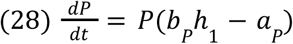

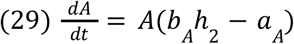

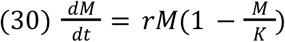

Here *M* is the tumor mass, η is the production rate of *h*_2_ secreted per one unit mass of tumor *M* and 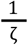 is the half-maximal inhibitory concentration of *h*_3_ on η *M*. Other parameters are as above.

We normalized the equations by substituting 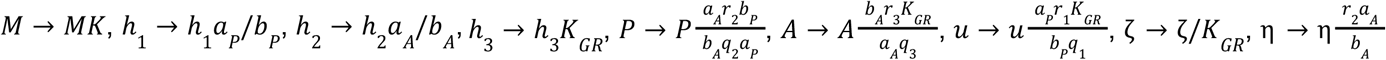 to obtain:

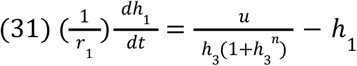

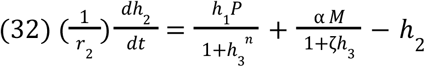

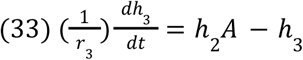

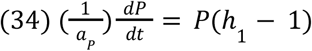

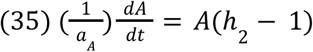

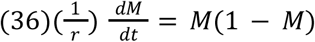

where we define α = η *K* as the rate of ACTH secretion from the tumor at its carrying capacity, which is the maximal ACTH secretion rate.

Steady state solutions below the bifurcation (α < 1 + ζ*A*, where *A* is defined implicitly by *u* = *A* + *A*^*n*+1^, and where native pituitary functional mass *P* > 0 shrinks to compensate for the tumor, and the tumor reaches its carrying capacity *M* = *K*):

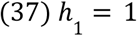

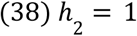

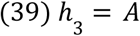

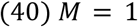

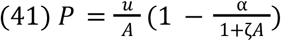

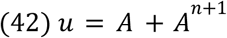

Steady state solutions above the bifurcation (α > 1 + ζ*A*, where *A* is defined implicitly by *u* = *A* + *A*^*n*+1^, native pituitary functional mass = 0, tumor in its carrying capacity *M* = *K*):

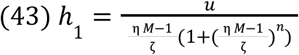

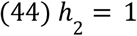

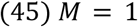

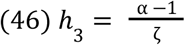

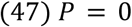

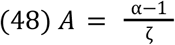

### Model equations for comparison of alternative HPA designs

To model alternative HPA designs, we started from the following equations from Karin et al.(21) for the full HPA axis:

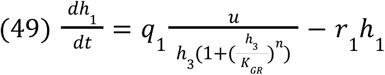

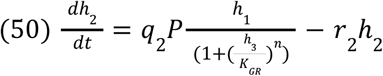

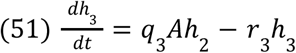

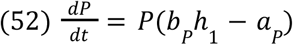

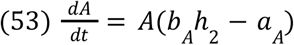

For the alternate circuit with a constant functional mass of the pituitary, we set *dP*/*dt* = 0 and *P* = *P*_0_, dropping Equation 52. For the circuit without a pituitary (equivalent to class 4 circuit design), we dropped Equations 50 and 52, and we replaced *h*_1_ with *h*_2_ in Equation 51.

